# Roles for stress response and cell wall biosynthesis pathways in caspofungin tolerance in *Cryptococcus neoformans*

**DOI:** 10.1101/465260

**Authors:** Kaila M. Pianalto, R. Blake Billmyre, Calla L. Telzrow, J. Andrew Alspaugh

**Author notes:** Publicly Available Data: Raw sequencing read data is available at the Sequence Read Archive through NCBI, accession number PRJNA501913. Current address: Stowers Institute for Medical Research, Kansas City, MO, 64110. **Corresponding Author**: J. Andrew Alspaugh, Box 102359, Duke University School of Medicine, 26 Durham, NC 27710.

## Abstract

Limited antifungal diversity and availability are growing problems for the treatment of fungal infections in the face of increasing drug resistance. The echinocandins, one of the newest classes of antifungal drugs, inhibit production of a crucial cell wall component. However, these compounds do not effectively inhibit the growth of the opportunistic fungal pathogen *Cryptococcus neoformans*, despite potent inhibition of the target enzyme. We therefore performed a forward genetic screen to identify cellular processes that mediate the relative tolerance of this organism to the echinocandin drug, caspofungin. Through these studies, we identified 14 genetic mutants that enhance caspofungin antifungal activity. Rather than directly affecting caspofungin antifungal activity, these mutations seem to prevent the activation of various stress-induced compensatory cellular processes. For example, the *pfa4*Δ mutant has defects in the palmitoylation and localization of many of its target proteins, including the Ras GTPase and the Chs3 chitin synthase which are both required for caspofungin tolerance. Similarly, we have confirmed the link between caspofungin treatment and calcineurin signaling in this organism, but we suggest a deeper mechanism in which caspofungin tolerance is mediated by multiple pathways downstream of calcineurin function. Additionally, a partial loss-of-function mutant of a COP9 signalosome component results in a highly caspofungin-susceptible strain of *C. neoformans*. In summary, we describe here several pathways in *C. neoformans* that contribute to the complex caspofungin tolerance phenotype in this organism.

## INTRODUCTION

Invasive fungal diseases primarily affect people with immune system defects, resulting in significant morbidity and mortality in these vulnerable patient populations (Park *et al*. 2009; Pyrgos *et al*. 2013; Rajasingham *et al*. 2017). Major challenges for effective treatment of systemic fungal infections include limited therapeutic options and availability, particularly in regions where fungal infection rates are highest (Loyse *et al*. 2013; Perfect and Bicanic 2014). Historically, it has been difficult to identify novel antifungal agents that are not also toxic to humans, since many cellular processes are highly conserved between humans and fungi. In the search for novel antifungal drugs, identification of fungal-specific cellular processes has been a major focus. The fungal cell wall represents a key structure for fungal viability, growth, and host evasion (Latgé 2007; Doering 2009; Gow and Hube 2012; O’Meara *et al*. 2013; Esher *et al*. 2018). Thus, compounds that target the production and maintenance of the fungal cell wall, with little to no effect on the human host, would make exciting and specific antifungal agents.

Echinocandins are cell wall-targeting antifungal compounds that have been identified and synthesized from natural products (Denning 2003; Letscher-Bru and Herbrecht 2003). These compounds inhibit the synthesis of β-1,3-glucan, a crucial cell wall component for many fungi (Taft *et al*. 1988; Kurtz *et al*. 1994; Kurtz and Douglas 1997; Maligie and Selitrennikoff 2005). Echinocandin antifungals, such as caspofungin, micafungin, and anidulafungin, are used extensively in clinical settings for the treatment of infections caused by diverse fungi. However, echinocandins, such as caspofungin, do not have potent antifungal activity against the fungal pathogen *Cryptococcus neoformans*, whose growth is only inhibited at high levels of caspofungin that are not clinically achievable in patients (Abruzzo *et al*. 1997; Bartizal *et al*. 1997; Espinel-Ingroff 1998). The fact that this drug is so ineffective against this fungus is surprising for a number of reasons. First, the gene that encodes the β-1,3-glucan synthase catalytic subunit in *C. neoformans*, *FKS1*, is essential in this organism (Thompson *et al*. 1999). Additionally, the *C. neoformans* enzyme is highly sensitive to caspofungin *in vitro*, even potentially at lower concentrations than species that are clinically susceptible to these drugs, such as *Aspergillus* species (Maligie and Selitrennikoff 2005). Based on these data, caspofungin could be expected to be an effective inhibitor of *C. neoformans* growth.

Given these observations, several investigators have tried to explain the discrepancy between the high sensitivity of the target enzyme activity and the high tolerance of the organism to caspofungin. Recent work using a fluorescently-tagged form of caspofungin has suggested that intracellular concentrations of caspofungin are low in wild-type *C. neoformans* (Huang *et al*. 2016). However, this could be due to either poor entry of caspofungin into the cell or rapid efflux or degradation of the drug. For example, the cell wall or the polysaccharide capsule could prevent accessibility of caspofungin, a high molecular-weight drug, to its target enzyme. Alternatively, caspofungin could be entering the cell but be rapidly eliminated from the cell through the action of a multi-drug resistance pump. Given the essentiality of the β-1,3-glucan synthase gene, true inhibition of this enzyme should result in detrimental effects to the cell. We hypothesize that *C. neoformans* expresses cellular factors that are important for its paradoxical tolerance to caspofungin. We predict that these processes allow *C. neoformans* to survive in the presence of caspofungin, leading to ineffective drug treatment.

In this work, we screened through two targeted deletion collections to identify *C. neoformans* mutants that were hypersensitive to caspofungin relative to the wild-type strain. In this way, we identified novel processes and pathways that facilitate echinocandin resistance in this organism. Here, we describe 14 genes and others in associated pathways that were identified in this screen and their roles in caspofungin tolerance. We also demonstrate possible mechanisms for how the cellular processes controlled by these genes might affect echinocandin resistance in *C. neoformans*.

## MATERIALS AND METHODS

### Strains, media, and growth conditions

The collections used for the caspofungin sensitivity screen consist of both the 2008 Madhani and 2015 Madhani plate collections, which were purchased from the Fungal Genetics Stock Center (Liu *et al*. 2008; Chun and Madhani 2010). The wild-type (WT) strain used in this study is the clinical strain H99 (Perfect *et al*. 1980). Newly generated strains, as well as strains from alternate sources, are listed in Table S1. A similar screen using a subset of these mutant strains was also recently performed (Huang *et al*. 2016).

Strains were maintained on yeast extract-peptone-dextrose (YPD) agar (10% yeast extract, 20% peptone, 2% dextrose, and 20% Bacto agar), and overnight cultures were incubated in YPD liquid medium. Drug susceptibility testing was performed in Yeast Nitrogen Base medium (1X YNB + 2% glucose) (Pfaller *et al*. 1990; Jessup *et al*. 1998). Cultures for microscopy were prepared in synthetic complete (SC) medium (1X YNB + 1X complete amino acids + 2% glucose).

### Strain creation

To generate the new and independent mutant strains used in this study, targeted gene deletion constructs were designed to replace the entire open reading frame with the neomycin (NEO) or nourseothricin (NAT) dominant selectable markers. Each knockout construct was generated using PCR overlap-extension and split selectable marker as described previously (Davidson *et al*. 2002; Kim *et al*. 2012). All constructs were transformed into the *C. neoformans* H99 strain by biolistic transformation as previously described (Toffaletti *et al*. 1993). All deletion primers used in this study can be found in Table S2A.

Upon transformation, strains were selected on YPD medium containing either nourseothricin or neomycin. Deletion mutants were checked by a combination of positive and negative confirmation PCRs demonstrating replacement of the WT locus with the mutant allele, followed by Southern blot to confirm single integration of the deletion constructs (data not shown).

Plasmids used in this study can be found in Table S3. Cloning primers can be found in Table S2B. The *MSH1* complementation plasmid was engineered by cloning the *MSH1* gene into the multiple cloning site of the pSDMA57 Safe Haven *NEO* plasmid (Arras *et al*. 2015). The COP9 complementation construct was generated by cloning the COP9 gene with its endogenous promoter and terminator into the pJAF1 plasmid (Fraser *et al*. 2003). The overexpressed *GFP*-*RHO1* construct was generated by cloning the *RHO1* gene plus its native terminator into the *BamHI* site of the pCN19 vector, which contains the histone H3 promoter and GFP with the *NAT* selection marker. The endogenous *FKS1*-*GFP* construct was engineered by cloning the following into the pUC19 vector: the *FKS1* gene (without promoter), the *GFP* gene, the *FKS1* terminator, and the *NEO* marker flanked by genomic sequence to target this construct to the *FKS1* locus. These plasmids were transformed as described and selected on neomycin or nourseothricin, and transformants were confirmed by a positive PCR to document the presence of the introduced allele. In the case of the *eFKS1*-*GFP* construct, PCRs to confirm integration of the construct into the endogenous *FKS1* locus were performed.

### Caspofungin sensitivity primary screen

A small pilot screen revealed that the calcineurin B subunit mutant was hypersensitive to caspofungin compared to the WT, agreeing with published data (Del Poeta *et al*. 2000). This strain became the standard against which to measure other potentially caspofungin-susceptible strains. To determine the optimal conditions under which to perform the caspofungin sensitivity screen, we assessed WT vs. *cnb1*Δ growth in YNB medium at caspofungin concentrations between 5 and 50 μg/mL with shaking at 150 rpm at 30° (Cancidas, Merck). We identified 15 μg/mL caspofungin to be a concentration at which the *cnb1*Δ mutant strain was markedly impaired for growth and the WT grew robustly. We then screened the strain collections of 3880 isolates, first pre-incubating in YPD liquid medium for 16 hours with shaking (150 rpm) at 30°. Cultures were diluted 1:10 in 96-well plates containing either YNB or YNB + caspofungin (15 μg/mL). Strains were incubated with shaking (150 rpm) at 30° for 24 hours, and growth was assessed by measuring OD_600_ on a FLUOStar Optima plate reader (BMG Labtech). Plates were also pin replicated to YPD plates to assess strain viability after incubation with caspofungin.

After screening, strains were divided into four groups based on caspofungin susceptibility: 1. Strains that were inviable post-caspofungin treatment; 2. Strains that were viable after caspofungin treatment, but had significantly decreased growth (OD_600_ that is greater than two standard deviations less than average WT OD_600_); 3. Strains that did not have significantly different growth from WT in caspofungin medium; and 4. Strains that did not grow in YNB.

### Disc-diffusion secondary screen

To confirm the caspofungin sensitivity of the mutants identified in the above screen, each mutant was incubated overnight in 150 uL of YPD in a 96-well plate with shaking, along with the WT and *cna1*Δ mutant strains as controls. Strains were diluted 1:200 in PBS, then 75 μL per strain was spread onto YNB agar in 6-well plates. Sterile filter discs were placed in the center of each well, and 5 μL of 7.5 mg/mL caspofungin was added to each disc. Plates were incubated at 30° for three days. After incubation, plates were imaged and zones of inhibition were measured. Mutant phenotypes were classified as WT-like, *cna1*Δ-like, or intermediate.

### Minimal Inhibitory Concentration (MIC) assay

MIC assays were performed according to modified CLSI standard methods for broth microdilution testing of antifungal susceptibility (Pfaller *et al*. 1990; CLSI 2008). In brief, cells were diluted in phosphate-buffered saline (PBS) to an OD_600_ of 0.25, then diluted 1:100 in YNB medium. Caspofungin was diluted in PBS., 2X working stocks of caspofungin were prepared in YNB medium, then caspofungin was serially diluted two-fold in 100 μL YNB medium. 100 μL of 1:100 dilution of cells was added to diluted drug in 96-well plates. Final concentration range of caspofungin was 200 to 0.39 μg/mL. Plates were incubated for 48 hours without shaking at 30° or 35°. After 48 hours, OD_600_ was measured on a FLUOStar Optima plate reader. MIC_50_ values were calculated by calculating relative growth using (drug-treated OD_600_/untreated OD_600_), with MIC_50_ corresponding to at least 50% decrease in relative growth.

### Checkerboard assay

Checkerboard assays to assess antifungal drug synergy using the Fractional Inhibitory Concentration index (FIC) for combinations of compounds were performed as described (NCCLS 1992; Franzot and Casadevall 1997). In brief, the WT strain H99 was inoculated from plated colonies into PBS at an OD_600_ of 0.25, and subsequently diluted 1:100 in YNB medium (or RPMI for clorgyline). Nikkomycin Z and tipifarnib stocks were diluted in PBS (Nikkomycin Z, Sigma-Aldrich; tipifarnib, Sigma-Aldrich). Manumycin A, clorgyline, and 2-bromopalmitate (2BP) were diluted in DMSO (Sigma-Aldrich). Caspofungin was diluted for a final concentration range between 100 μg/mL and 1.5625 μg/mL, and the test drugs were diluted to the following final concentration ranges: nikkomycin Z 400 to 0.78125 μg/mL, tipifarnib 400 to 0.78125 μg/mL, manumycin A 40 to 0.078 μM, 2BP 400 to 0.78125 μM, and clorgyline 100 to 1.5625 μM. Assays were incubated at 30° for nikkomycin Z and 37° for clorgyline, manumycin A, tipifarnib, and 2BP. FIC index values for a combination of compounds A and B were calculated as:

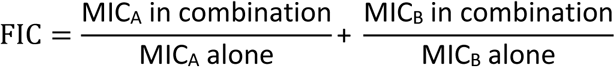

where an FIC index value of <1.0 is considered synergistic (with <0.5 considered strongly synergistic), additive if the value was 1.0, autonomous if the value was between 1.0 and 2.0, and antagonistic if the FIC index was greater than 2.0 (Franzot and Casadevall 1997).

### Chitin and chitosan assay

Chitin and chitosan content of *C. neoformans* cell walls was assessed as described (Banks *et al*. 2005). Briefly, cells were incubated overnight in YPD. Cultured cells were then diluted to an OD of 0.8 and cultured in SC or SC + 15 μg/mL caspofungin for 6 hours. Cells were divided and lyophilized, then either mock treated or treated with acetic anhydride to acetylate chitosan to form chitin. Cell walls were then digested with 5 mg/mL chitinase for 72 hours. GlcNAc monomer levels were assessed by a DMAB (*p*-dimethylaminobenzaldehyde) colorimetric assay and read on a FLUOStar Optima plate reader. Acetic anhydride samples represented levels of both chitin and chitosan in the cell wall, while untreated samples represented chitin alone. Chitosan levels were calculated as the difference between the acetic anhydride-treated and the untreated samples. Data was analyzed using a two-way ANOVA, followed by t-tests to determine statistical significance.

### Cell Wall Staining and Microscopy

Cells were prepared for cell wall staining as described (Ost *et al*. 2017). WT cells were cultured overnight in YPD medium. Overnight cultures were diluted to an OD_600_ of 1 in 15 mL SC or SC plus caspofungin (5, 10 or 20 μg/mL caspofungin). At indicated timepoints, 1 mL aliquots of each culture were collected and stained with calcofluor white (CFW). Cells were pelleted at 5000 rpm for two minutes, then resuspended in 100 μL PBS + 25 μg/mL CFW and incubated in the dark at room temperature for 10 minutes. Cells were then washed two times with PBS and resuspended in 50 μL PBS for imaging. Strains were imaged on a Zeiss Axio Imager A1 fluorescence microscope equipped with an Axio-Cam MRM digital camera to capture both DIC and fluorescent images. Cell wall staining fluorescent intensity was analyzed using Fiji software, and the mean gray values were analyzed (Schindelin *et al*. 2012). Data presented represents the average fluorescence values. Data was analyzed using a two-way ANOVA, followed by t-tests to determine statistical significance.

For fluorescent fusion protein microscopy, strains were incubated overnight in SC medium. Overnight cultures were pelleted at 3000 rpm for five minutes and resuspended in SC or SC + 15 μg/mL caspofungin. Strains were incubated for 90 minutes, with aliquots collected for imaging at 15-minute intervals. Aliquots were incubated with NucBlue Live Ready Probes reagent for five minutes, pelleted at 5000 rpm for two minutes, then resuspended in 50 μL SC (Thermo Fisher Scientific). Strains were imaged on a Zeiss Axio Imager A1 fluorescence microscope equipped with an Axio-Cam MRM digital camera to capture both DIC and fluorescent images. For the Fks1-GFP and GFP-Rho1 localization experiments, overnight cultures were normalized to an OD_600_ of 2.0 in SC plus 0, 5, 10, or 15 μg/mL caspofungin and imaged at 20-minute intervals for 1.5 hours.

### Whole Genome Sequencing, Alignment, and Variant Calling

Whole-genome sequencing was performed on both the WT background strain and the *msh1*Δ mutant strain by the Duke Center for Genome and Computational Biology Genome Sequencing Shared Resource using an Illumina MiSeq instrument. Paired end libraries were sequenced with read lengths of 251 bases. Reads were aligned to the version 3 H99 genome (Janbon *et al*. 2014) using BWA-MEM with default settings (Li and Durbin 2009). The GATK best practices pipeline (McKenna *et al*. 2010) was used in combination with SAMtools (Li *et al*. 2009) and Picard to realign reads before SNP calling using the UnifiedGenotyper component of GATK with the haploid ploidy setting. The resulting VCFs were filtered using VCFtools (Danecek *et al*. 2011) and annotated for variant effect using SnpEff (Cingolani *et al*. 2012). Heterozygous calls were removed as presumed mismapped repetitive regions. Raw reads are available on the NCBI Sequence Read Archive under accession number PRJNA501913.

### RNA Preparation and Quantitative Real-Time PCR

The WT strain was grown in YPD overnight at 30° with shaking. Cells were then inoculated at an OD_600_ of 1.5 into 5 mL SC, SC + 10 μg/mL caspofungin, or SC + 15 μg/mL caspofungin. Cultures were incubated at 30° with shaking for 90 minutes, then cells were harvested by centrifugation at 3000 rpm for five minutes and lyophilized. RNA was isolated using a RNeasy Plant Mini Kit (Qiagen), with the addition of bead beating for one minute prior to lysis and on-column DNase treatment (Qiagen). cDNA was prepared using the AffinityScript QPCR cDNA synthesis kit using oligo-dT primers to bias for mRNA transcripts (Agilent Genomics). Quantitative Real-Time PCR was performed using PowerUp SYBR Green Master mix (Applied Biosystems) on a QuantStudio 6 Flex system. Real-time PCR primers are listed in Table S2C (Esher *et al*. 2018). Data was analyzed using a two-way ANOVA, followed by t-tests to determine statistical significance.

### Reagent and Data Availability

Strains and plasmids are available upon request. File S1 contains a list and descriptions of all supplemental files. Sequence data are available at the NCBI Sequence Read Archive under the accession number PRJNA501913. Supplemental files have been submitted to figshare.

## RESULTS

### Initial screen for processes contributing to caspofungin tolerance in *Cryptococcus neoformans*

We performed a forward genetic screen of targeted deletion mutants to identify cellular processes that contribute to *C. neoformans* tolerance to caspofungin treatment. Using two screening methods in sequence, we screened 3,880 mutants for altered growth during caspofungin treatment (Liu *et al*. 2008; Chun and Madhani 2010). Our initial sensitive but qualitative primary screen for altered caspofungin susceptibility identified 232 mutants with reduced caspofungin tolerance compared to WT. These strains were subsequently tested in two secondary screens for caspofungin susceptibility using both disc-diffusion and broth microdilution assays to compare caspofungin susceptibility to the WT strain, as well as to a *cna1*Δ strain with a mutation in the calcineurin A subunit gene. Strains with altered calcineurin function are known to be more susceptible to caspofungin (Del Poeta *et al*. 2000). Of the mutants identified in the primary screen, 14 were confirmed to have caspofungin susceptibility similar to or greater than the *cna1*Δ mutant strain (Table 1). The remaining strains displayed only minimal increases in sensitivity to caspofungin, and they were not tested further.

**Table 1.** Caspofungin MICs for Screen Hits

| Gene locus tag | Gene product mutated | MIC, µg/mL |
| --- | --- | --- |
| CNAG_00375 | SAGA complex histone acetyltransferase (Gcn5) | 12.5 |
| CNAG_02682 | hypothetical protein (Msh1 homolog) | 0.78 |
| CNAG_05070 | sulfite reductase (NADPH) hemoprotein, beta-component | 0.78 |
| CNAG_06902 | hypothetical protein | 12.5 |
| CNAG_02236 | Type 2A-like serine/threonine-protein phosphatase (Ppg1) | 0.78 |
| CNAG_03981 | palmitoyltransferase Pfa4 | 6.25 |
| CNAG_03841 | hypothetical protein | 12.5 |
| CNAG_02292 | copper chaperone Lys7 | 12.5 |
| CNAG_04992 | hypothetical protein | 12.5 |
| CNAG_03080 | fatty acid elongase | 12.5 |
| CNAG_07636 | chitin synthase regulator (Csr2) | 3.125 |
| CNAG_02891 | endoplasmic reticulum rhodanese-like protein (Rdl2) | 12.5 |
| CNAG_01717 | cell differentiation protein Rcd1 | 12.5 |
| CNAG_00888 | Calcineurin B subunit | 3.125 |

Within the list of sensitive mutants, we identified multiple biological processes that seem to be important for caspofungin tolerance based on gene ontology analysis (full GO analysis Table S4) (Stajich *et al*. 2012). Multiple genes that have been associated with responses to stress were identified, including those encoding the gene products chitin synthase regulator 2, Ppg1 phosphatase, the Gcn5 histone acetyltransferase, and the Msh1 mismatch repair protein (Banks *et al*. 2005; Gerik *et al*. 2005; O’Meara *et al*. 2010; Boyce *et al*. 2017). Additionally, we identified two genes involved in cell integrity, *CSR1* and *PPG1*. We also found multiple proteins that have potential antioxidant roles, such as an Rdl2 Rhodanese homolog and a putative Lys7 homolog which typically partners with superoxide dismutase 1 (Culotta *et al*. 1997; Orozco *et al*. 2012). A list of mutants with the most striking MIC values are listed in Table 1.

### Calcineurin signaling plays a role in caspofungin tolerance in *C. neoformans*

In previous *in vitro* studies, the calcineurin inhibitor FK506 has demonstrated synergistic interactions with caspofungin against *C. neoformans* (Del Poeta *et al*. 2000). In our caspofungin sensitivity screen, we identified a mutant of the calcineurin B regulatory subunit, *cnb1*Δ, to be highly sensitive to caspofungin. Indeed, mutants of both the calcineurin A catalytic and calcineurin B regulatory subunits—*cna1*Δ and *cnb1*Δ, respectively—exhibit an eight-fold increase in caspofungin sensitivity when compared to the WT strain (Table 1). Identification of this mutant with a defect in calcineurin signaling, a pathway that is known to be involved with caspofungin tolerance, largely validated the screening approaches used in this study.

To determine how calcineurin signaling might be mediating caspofungin tolerance in *C. neoformans*, we assessed caspofungin sensitivity for mutants of known targets of calcineurin phosphatase activity. The Crz1 transcription factor is activated by the calcineurin protein, and Crz1 mediates many of the known effects of this calcineurin signaling. Accordingly, we assessed whether an mCherry-tagged Crz1 fusion protein (Crz1-mCherry) localizes to the nucleus after treatment with caspofungin (Chow *et al*. 2017). We assessed Crz1 nuclear localization by examining Crz1-mCherry co-localization with the GFP-Nop1 nucleolar marker as well as with DAPI staining. In untreated cells, Crz1-mCherry remains localized in the cytosol. In 15 μg/mL of caspofungin, we determined that Crz1-mCherry strongly localizes to the nucleus after 45 minutes of incubation (Figure 1). The nuclear localization of Crz1-mCherry suggests that Crz1 is likely being activated under these conditions. However, the *crz1*Δ mutant displayed a caspofungin MIC more similar to WT than to the *cna1*Δ or *cnb1*Δ strains (Table 2). Together, these results suggest that, although Crz1 is being activated during caspofungin treatment, a Crz1-independent target (or targets) of calcineurin activity is required for full caspofungin tolerance. These data are consistent with recent reports of both Crz1-dependent and -independent processes downstream of *C. neoformans* calcineurin signaling (Lev *et al*. 2012; Chow *et al*. 2017). To attempt to identify other calcineurin targets that might be involved in caspofungin tolerance, we assessed the caspofungin susceptibility of mutants of genes expressing several protein targets of calcineurin-mediated dephosphorylation (Table S5) (Park *et al*. 2016). However, none of those tested displayed increased caspofungin susceptibility, suggesting that other as yet unidentified targets mediate this phenomenon in *C. neoformans*.

**Pfa4 plays a role in caspofungin tolerance through regulation of its target proteins**.

**Figure 1.**
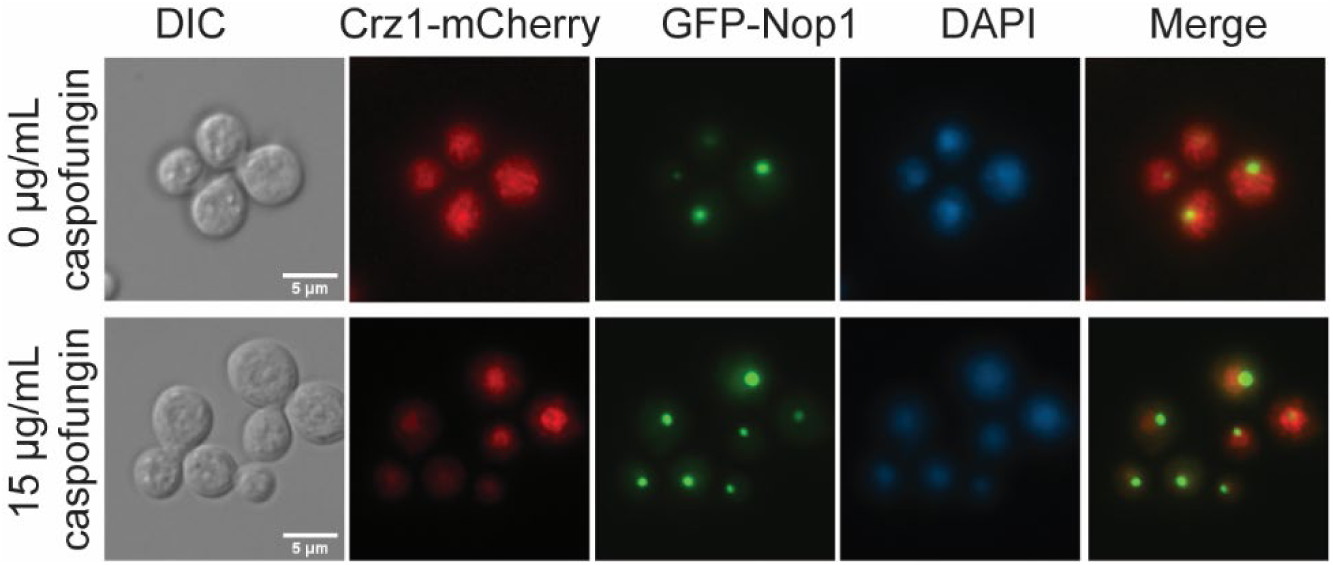
Crz1 is activated and localizes to the nucleus in response to caspofungin treatment. A strain expressing Crz1-mCherry and GFP-Nop1 (nucleolar marker) was incubated in SC with either 0 or 15 μg/mL caspofungin for 45 minutes, then stained with NucBlue Live Cell nuclear stain for 5 minutes and imaged on a Zeiss AxioVision epifluorescence microscope. Scale bars represent 5 microns.

**Table 2.**
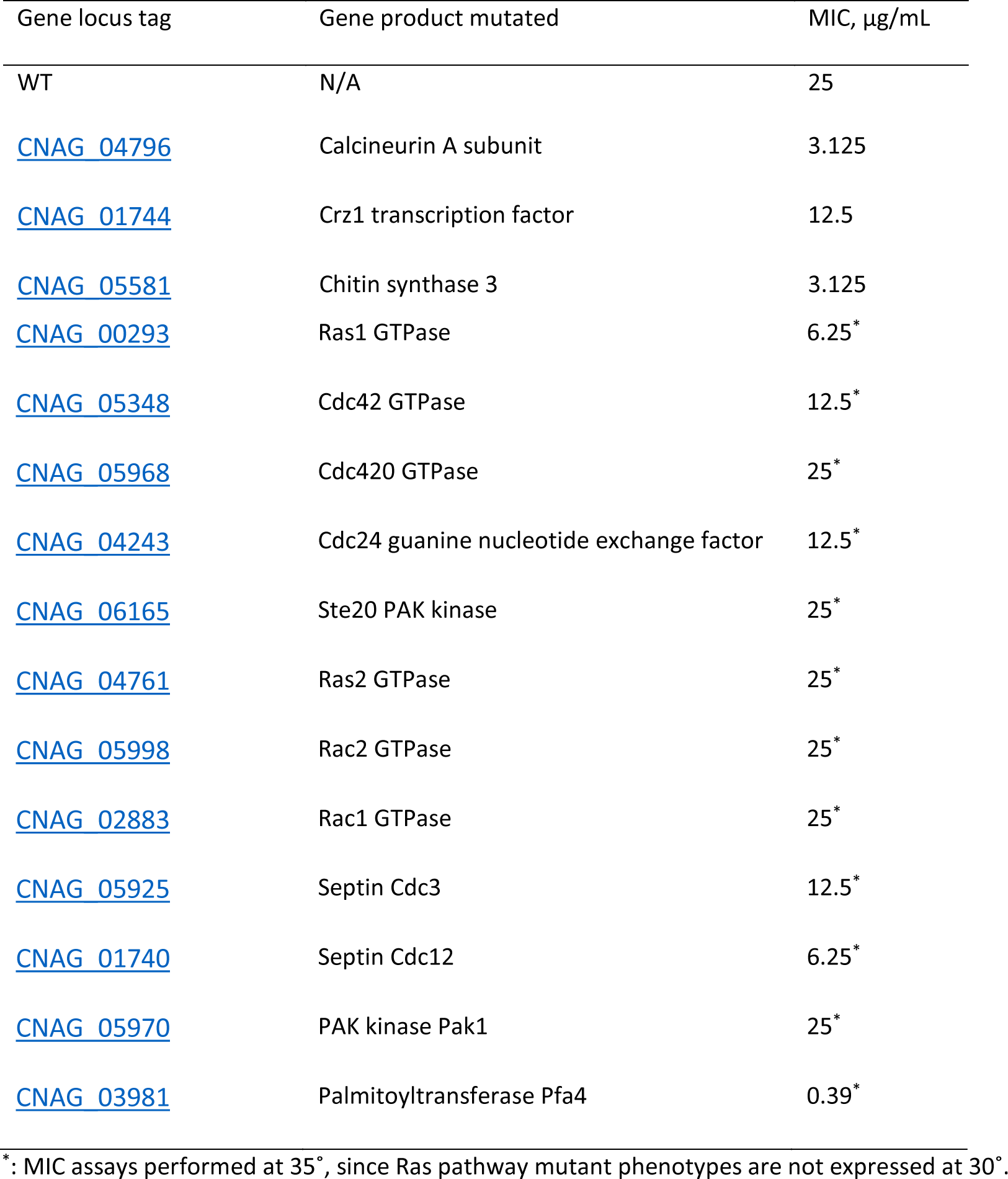
Additional Caspofungin MICs

| Gene locus tag | Gene product mutated | MIC, µg/mL |
| --- | --- | --- |
| WT | N/A | 25 |
| CNAG_04796 | Calcineurin A subunit | 3.125 |
| CNAG_01744 | Crz1 transcription factor | 12.5 |
| CNAG_05581 | Chitin synthase 3 | 3.125 |
| CNAG_00293 | Ras1 GTPase | 6.25* |
| CNAG_05348 | Cdc42 GTPase | 12.5* |
| CNAG_05968 | Cdc420 GTPase | 25* |
| CNAG_04243 | Cdc24 guanine nucleotide exchange factor | 12.5* |
| CNAG_06165 | Ste20 PAK kinase | 25* |
| CNAG_04761 | Ras2 GTPase | 25* |
| CNAG_05998 | Rac2 GTPase | 25* |
| CNAG_02883 | Rac1 GTPase | 25* |
| CNAG_05925 | Septin Cdc3 | 12.5* |
| CNAG_01740 | Septin Cdc12 | 6.25* |
| CNAG_05970 | PAK kinase Pak1 | 25* |
| CNAG_03981 | Palmitoyltransferase Pfa4 | 0.39* |

\*: MIC assays performed at 35°, since Ras pathway mutant phenotypes are not expressed at 30°.

### Pfa4 palmitoyltransferase mutant is hypersensitive to caspofungin

The Pfa4 palmitoyltransferase is required for the addition of palmitoyl groups to various *Cryptococcus neoformans* proteins (Nichols *et al*. 2015; Santiago-Tirado *et al*. 2015). This post-translational modification is necessary for the proper localization and function of these target proteins. In our MIC assays, we documented a four-fold increase in caspofungin susceptibility for the *pfa4*Δ strain (Table 2). Since Pfa4 is responsible for the regulation of various functions within the cell, it is likely that this caspofungin sensitivity is due to dysregulation of one or more Pfa4 palmitoylation targets. Therefore, the caspofungin susceptibility for mutants in several Pfa4-regulated gene products was assessed.

Additionally, since Pfa4-mediated palmitoylation also seems to play a role in caspofungin tolerance in *C. neoformans*, we assessed whether there might be synergy between caspofungin and inhibitors of palmitoyltransferases. Though the competitive palmitoyltransferase inhibitor 2-bromopalmitate (2BP) displays some activity against *Aspergillus fumigatus*, we found that it displayed poor activity against *C. neoformans* (Jennings *et al*. 2008; Fortwendel *et al*. 2012). Accordingly, we found that 2BP and caspofungin displayed completely autonomous activity, with an FIC index of 2 for these two compounds (Table 3).

**Table 3.**
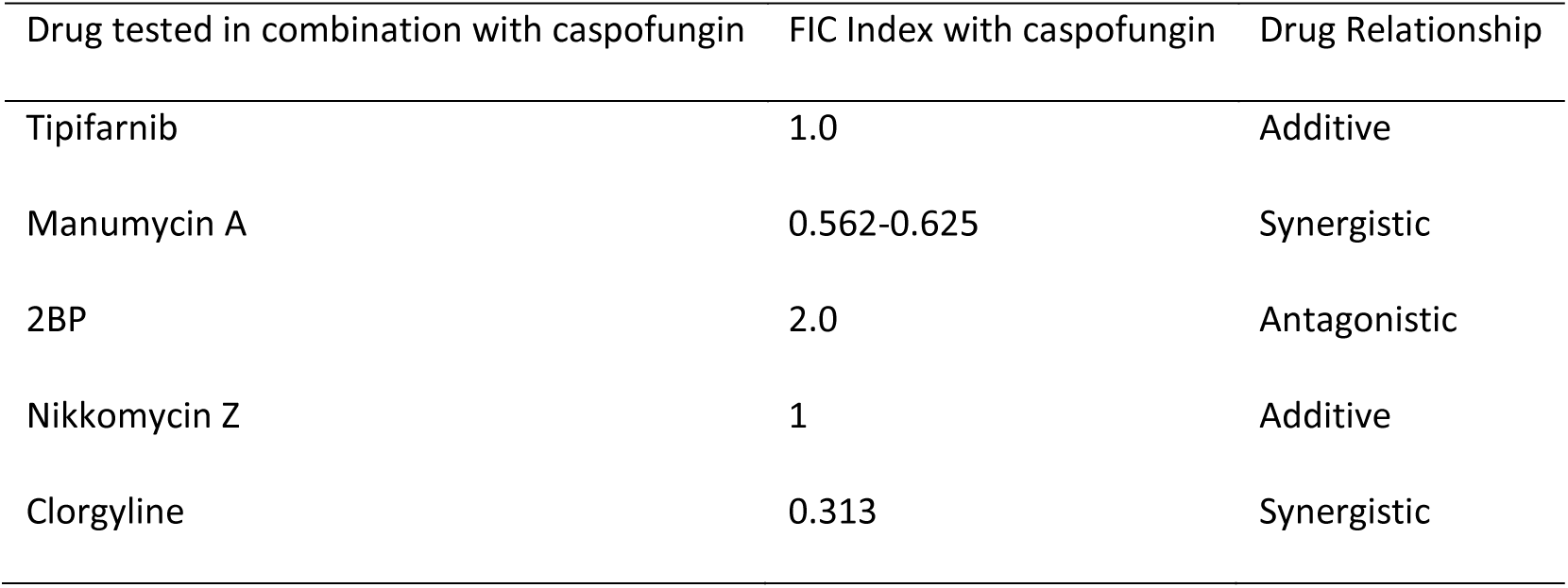
Combination Fractional Inhibitory Concentration (FIC) Indices

| Drug tested in combination with caspofungin | FIC Index with caspofungin | Drug Relationship |
| --- | --- | --- |
| Tipifarnib | 1.0 | Additive |
| Manumycin A | 0.562-0.625 | Synergistic |
| 2BP | 2.0 | Antagonistic |
| Nikkomycin Z | 1 | Additive |
| Clorgyline | 0.313 | Synergistic |

### Ras signaling

Pfa4 palmitoylates the *C. neoformans* Ras1 GTPase, and this post-translational modification is required for proper subcellular localization of Ras1 as well as its function (Nichols *et al*. 2015). Since Ras1 is required for thermotolerance of *C. neoformans*, the *pfa4*Δ mutant is accordingly growth defective at elevated temperatures. To determine whether the caspofungin susceptibility of the *pfa4*Δ mutant is reflected in this downstream target pathway, we assessed caspofungin susceptibility for the *ras1*Δ mutant, as well as for mutants in the Ras1 morphogenesis pathway (mediated by Rac proteins), and the Ras1 thermotolerance pathway (mediated by Cdc24/Cdc42 and the septin proteins) (Figure 2) (Waugh *et al*. 2002; Nichols *et al*. 2007; Ballou *et al*. 2009, 2013a; b). The *ras1*Δ mutant is four-fold more sensitive to caspofungin than WT. Increases in caspofungin susceptibility were also noted for the *cdc42A* and *cdc24Δ* mutants, as well as the *cdc3*Δ and *cdc12*Δ septin mutants, which are further downstream effectors of the *C. neoformans* Ras thermotolerance pathway (Nichols *et al*. 2007; Ballou *et al*. 2009, 2013a). These proteins mediate dynamic actin cytoskeletal changes required for budding and cell division in response to cell stresses such as elevated temperature. In contrast, the Rac1 and Rac2 proteins are not required for caspofungin tolerance; these Ras1-mediated GTPases are involved in a distinct signaling pathway controlling morphological transitions, such as hyphal formation during mating (Ballou *et al*. 2013b). Therefore, The Pfa4-Ras1-Cdc42-septin protein pathway seems to be required to optimally support *C. neoformans* growth in the presence of echinocandins, and inhibition of this signaling axis results in notable increases in caspofungin susceptibility.

**Figure 2.**
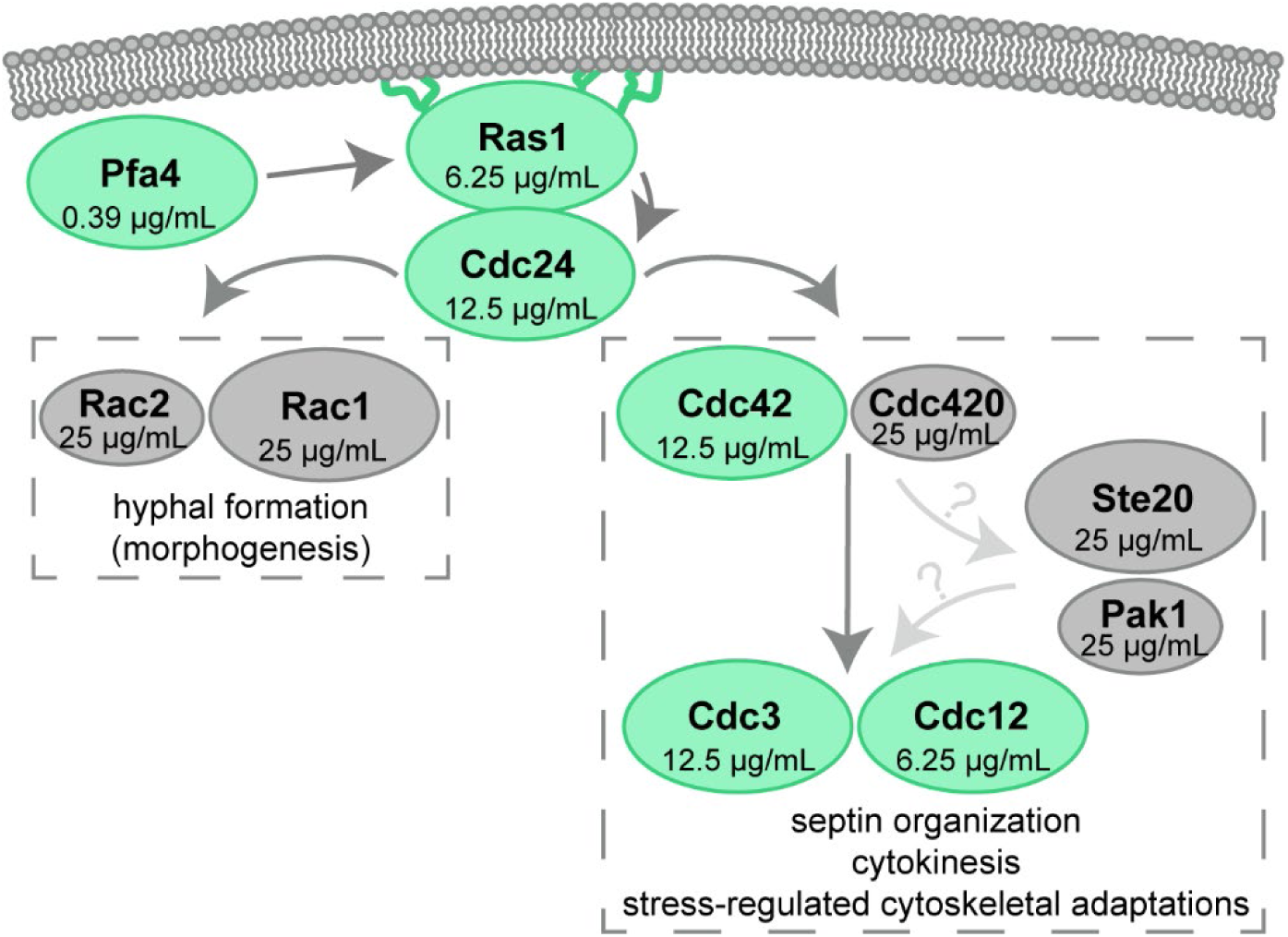
Ras1 signaling partner mutants are differentially affected by caspofungin treatment. Model of Ras signaling in *C. neoformans*. Paired paralogs are represented as the major (larger oval) and minor (smaller oval) paralog (Ballou *et al*. 2013a). Caspofungin MICs at 35° for each component are presented within each oval. The green ovals represent mutants that displayed increased caspofungin susceptibility, while the grey ovals displayed WT caspofungin susceptibility.

Although specific inhibition of fungal Ras activity is limited by the highly conserved nature of this protein, recent investigations have explored inhibitors of Ras-modifying enzymes, such as farnesyltransferases, as antifungal agents (Hast *et al*. 2011; Selvig *et al*. 2013; Esher *et al*. 2016; Pianalto and Alspaugh 2016). For proper localization and function, Ras-like GTPases also require, in addition to palmitoylation, the post-translational addition of lipophilic prenyl groups by farnesyltransferases (FTases) to C-terminal cysteine residues. We therefore assessed synergy between caspofungin and two different protein-farnesyltransferase inhibitors (FTIs), tipifarnib and Manumycin A (Hast *et al*. 2011). Tipifarnib displayed additivity with caspofungin against *C. neoformans*, with an FIC index of 1 (Table 3). However, the FTI Manumycin A displayed synergy with caspofungin, with an FIC index of 0.56 (Franzot and Casadevall 1997). Given the limited intrinsic antifungal activity of these first-generation FTIs, these results suggest that Ras inhibitors with greater anti-cryptococcal activity might be promising co-administered agents to augment the effect of caspofungin against *C. neoformans*.

### Chitin synthase

One of the most prominent targets of Pfa4 palmitoyltransferase activity is the Chs3 chitin synthase, which is responsible for the biosynthesis of chitin that is destined to become chitosan in the cell wall (Banks *et al*. 2005; Baker *et al*. 2007). Pfa4 is required for the proper localization of the Chs3 chitin synthase to the cell surface, which is necessary for proper Chs3 function in cell wall biosynthesis and maintenance (Santiago-Tirado *et al*. 2015). In addition to the *pfa4*Δ mutant strain, our screen also identified a mutant of the chitin synthase regulator associated with Chs3 function, *csr2*Δ, which displayed a similar, though slightly more severe, caspofungin sensitivity phenotype (Table 2) (Banks *et al*. 2005). We therefore assessed the caspofungin susceptibility of a *chs3*Δ strain to determine if misregulation of this protein and lack of proper chitin and chitosan deposition might contribute to the caspofungin susceptibility of the *pfa4*Δ mutant strain. The *chs3*Δ mutant strain was eight-fold more sensitive to caspofungin than the WT strain (Table 2). These results are consistent with those in *Aspergillus fumigatus* and *Candida* species in which compensatory increases in cell wall chitin are induced upon exposure of these fungi to caspofungin (Walker *et al*. 2008; Fortwendel *et al*. 2010; Verwer *et al*. 2012; Walker *et al*. 2012). In fact, similar caspofungin-dependent increases in *C. neoformans* cell wall chitin content were observed using the chitin-binding dye, calcofluor white (CFW). A dose-dependent increase in CFW staining was observed for cells treated with caspofungin compared to untreated cells, with the most significant differences being at the highest concentration of caspofungin (Figure 3A). Accordingly, when we assessed cell wall chitin and chitosan composition using an *in vitro* colorimetric assay (Banks *et al*. 2005), we found that *C. neoformans* cells treated with 15 μg/mL caspofungin have an approximately 2.5-fold increase in both chitin and chitosan compared to untreated cells (Figure 3B).

**Figure 3.**
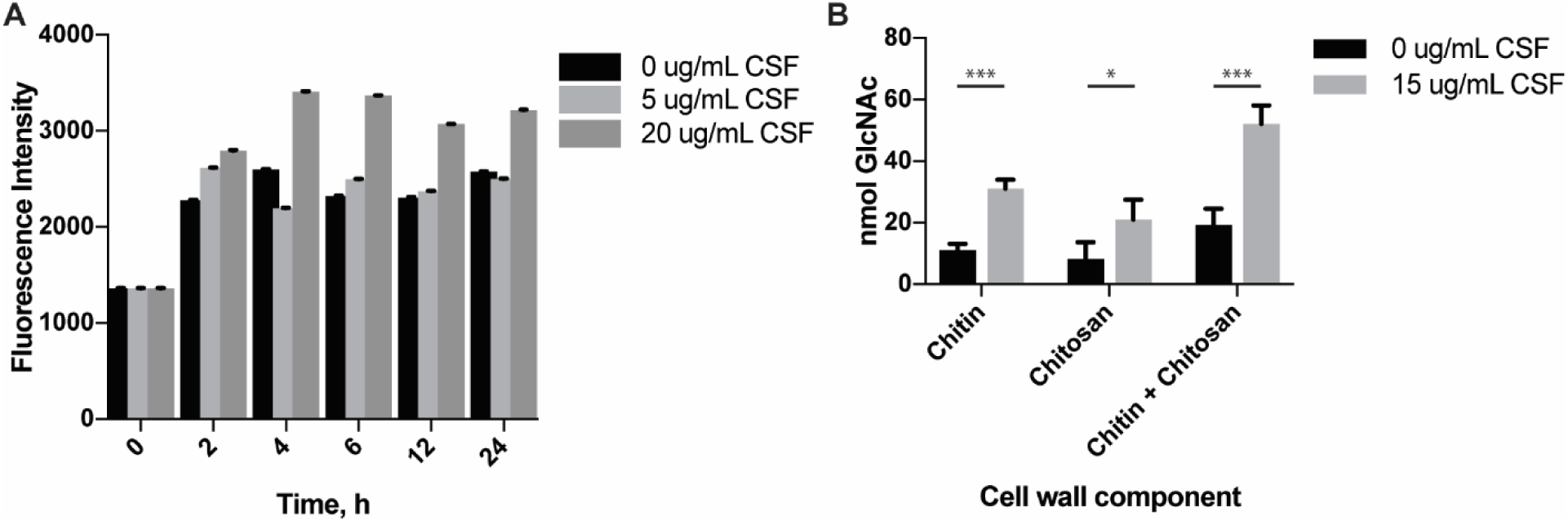
Cell wall chitin and chitosan levels increase during caspofungin treatment. **A**. Quantification of calcofluor white (CFW) staining of *C. neoformans* WT cells treated with caspofungin. Cells were incubated in SC medium with 0, 5, or 20 μg/mL caspofungin over 24 hours. At each timepoint, cells were harvested and stained with CFW to assess total cell wall chito-oligomer content, then imaged on a Zeiss AxioVision epifluorescence microscope. Images were masked for fluorescence, and average fluorescent intensity for each cell was quantified using ImageJ. Each timepoint represents quantification of greater than 80 cells over at least 3 images. Error bars represent SEM. CSF = caspofungin. **B**. Quantification of cell wall chitin and chitosan using DMAB colorimetric assay. Cells were incubated in SC +/- 15 μg/mL caspofungin for 6 hours, then harvested and assay performed as described. Error bars represent SEM. Statistics performed were 2-way ANOVA, followed by pair-wise t-tests. ^∗^, p<0.05; ^∗∗∗^, p<0.001.

Since the *chs3*Δ chitin synthase mutant strain is so susceptible to the effects of caspofungin, we hypothesized that the co-administration of caspofungin with the chitin synthase inhibitor Nikkomycin Z might result in a similar synergistic antifungal effect. We tested antifungal drug interactions using a checkerboard MIC assay, using varying concentrations of each drug in combination. In this assay, we did not observe synergy between caspofungin and Nikkomycin Z, one of the few known inhibitors of chitin synthesis in some fungi (Gaughran *et al*. 1994; Li and Rinaldi 1999). The FIC index, a marker of drug interaction, was 1.0, suggesting an additive effect between the two compounds rather than drug synergy (Table 3). However, historically, nikkomycin Z has not been an effective inhibitor of *C. neoformans* growth or chitin biosynthesis. Based on these genetic studies, there is potential for synergy between caspofungin and future, more effective chitin synthase inhibitors.

### A MutS homolog mutant is hypersensitive to caspofungin

One of the strains identified as being highly caspofungin-susceptible in the initial screen had a targeted mutation in an *MSH1* ortholog. Msh1 is a MutS homolog that is predicted to mediate DNA mismatch repair processes in the mitochondrial genome (Reenan and Kolodner 1992). This strain was highly susceptible to caspofungin (MIC 0.78 μg/mL). However, independent *msh1*Δ deletion strains were not hypersensitive to caspofungin, suggesting that another, silent mutation was responsible for this drug-sensitive phenotype in a strain with high mutagenic potential. Whole genome sequencing of this strain revealed three non-synonymous nucleotide polymorphisms in the genomic sequence. One of these mutations was a nonsense mutation (Glu313Stop) in the *CSN4* gene encoding subunit 4 of the constitutive photomorphogenesis (COP9) signalosome (*CSN4STOP* strain) (Table S6). Multiple attempts were unable to make a strain with a complete deletion of *CSN4*, suggesting that this gene is essential for viability under standard laboratory growth conditions. However, introduction of a full-length, WT *CSN4* gene into the *msh1*Δ (*csn4*STOP) mutant resulted in full complementation of the caspofungin hypersensitive phenotype. These combined results suggest that the *csn4*Δ mutation in this strain does not confer complete loss of function of this protein, and that a fully functional COP9 signalosome is required for caspofungin tolerance.

In addition to the nonsense mutation in the *CSN4* COP9 signalosome gene, multiple nonsynonymous mutations, including three separate frameshift mutations, were identified in the mitochondrial genome (Table S6). In fact, there was a thousand-fold difference in the number of mutations per base in the mitochondrial genome (3.7 × 10^−4^) versus the nuclear genome (3.7 × 10^−7^), consistent with recent data showing that *MSH1* does not affect the nuclear mutation rate (Litter *et al*. 2005; Janbon *et al*. 2014; Boyce *et al*. 2017). These data suggest that *C. neoformans MSH1* is likely acting as a mismatch repair protein for the mitochondrial genome.

### Cell wall gene expression is altered during caspofungin treatment

Since compensatory cell wall gene expression and activity, specifically chitin synthesis, has been demonstrated to play a role in paradoxical growth and resistance during caspofungin treatment, we assessed whether expression of cell wall biosynthesis genes was altered in response to caspofungin treatment. We assessed the transcript abundance for genes involved in the synthesis of chitin (*CHS1*, *CHS2*, *CHS3*, *CHS4*, *CHS5*, *CHS6*, *CHS7*, and *CHS8*), chitosan (*CDA1*, *CDA2*, and *CDA3*), α-1,3-glucan (*AGS1*), β-1,3-glucan (*FKS1*), and β-1,6-glucan (*KRE6* and *SKN7*) (Thompson *et al*. 1999; Banks *et al*. 2005; Reese *et al*. 2007; Gilbert *et al*. 2010; Esher *et al*. 2018). Many of these cell wall biosynthesis genes demonstrate altered regulation in response to caspofungin treatment (Figure 4). *CHS1*, *CHS2*, *CHS4*, *CHS7*, *SKN1*, *CDA1*, and *FKS1* demonstrated significantly increased expression, especially at the highest concentration of caspofungin tested in this experiment, representing a possible compensatory response. These results reflect some of the compensatory processes seen in other fungi, such as *Candida* species and *Aspergillus fumigatus*, in which increased cell wall chitin is a mechanism for these fungi to overcome the effects of caspofungin treatment (Walker *et al*. 2008; Fortwendel *et al*. 2010). However, we noted potential involvement of more diverse cell wall components. Additionally, the upregulation of *SKN1* suggests a potential role for β-1,6-glucan synthesis in the caspofungin tolerance mechanism for *C. neoformans*. In light of previous data showing that there is a decrease in β-1,6-glucan staining via immunoelectron microscopy, there is the potential that the upregulation of β-1,6-glucan biosynthetic genes could represent transcriptional compensation for secondary cell wall effects after drug treatment (Feldmesser *et al*. 2000). Interestingly, *CHS5*, *CHS6*, *CDA2*, and *CDA3* all displayed decreased expression during caspofungin treatment. Therefore, though not all cell wall-associated genes are upregulated in response to caspofungin, there seems to be a decisive alteration in the expression of cell wall biosynthesis and modification genes, reflecting a coordinated compensatory response to the cell wall inhibitor caspofungin.

**Figure 4.**
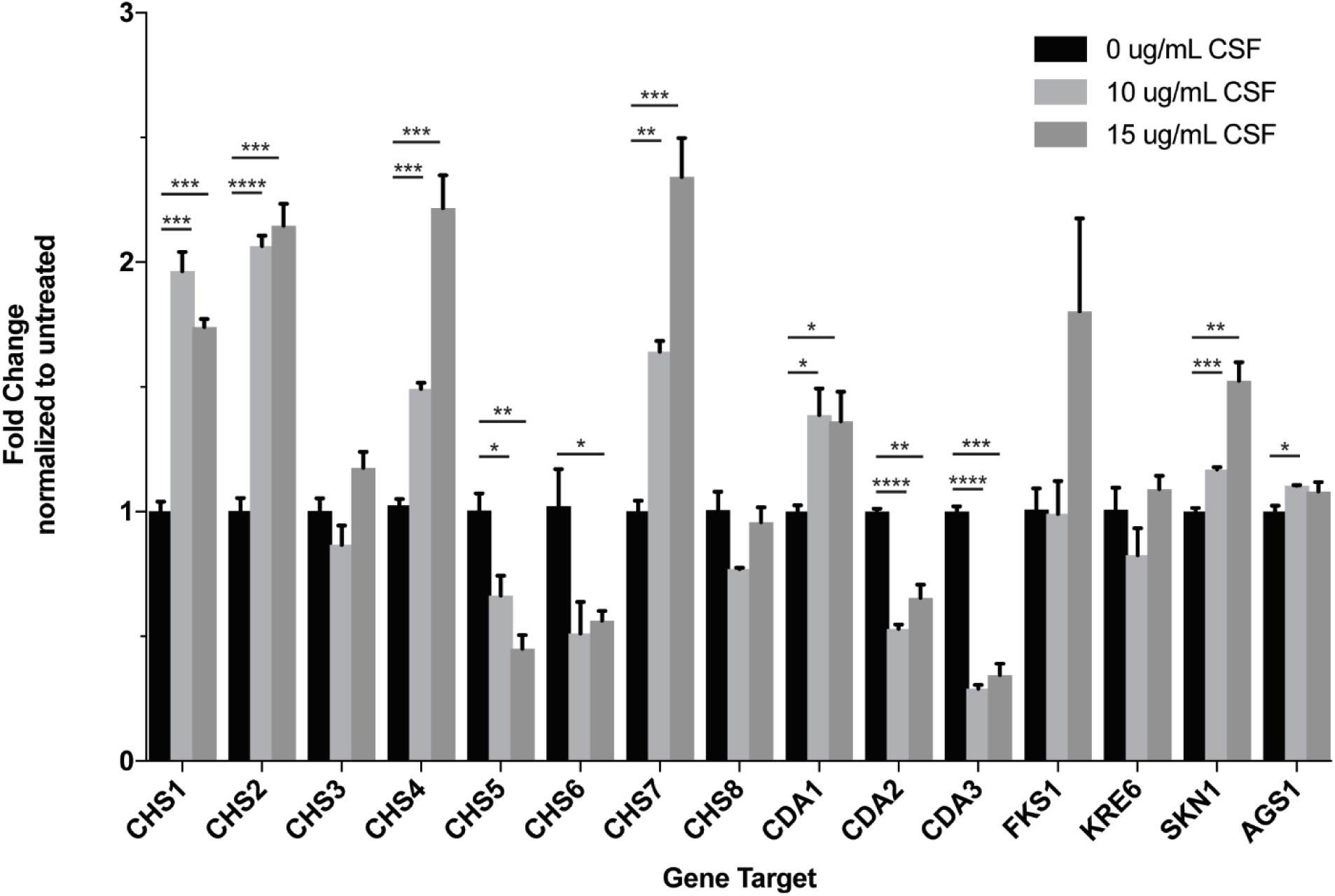
Cell wall biosynthesis gene expression is altered in response to caspofungin. WT *C. neoformans* cells were incubated in SC containing 0, 10, or 15 μg/mL caspofungin for 90 minutes, followed by RNA purification and quantitative real-time PCR for the indicated target genes, using the *GPD1* gene as an internal control. The results represent average values for biological triplicate samples. Statistical significance was assessed using 2-way ANOVA, followed by t-test for pairwise comparisons. ^∗^, p<0.05; ^∗∗^, p<0.01; ^∗∗∗^, p<0.001; ^∗∗∗∗^, p<0.0001. CSF = caspofungin.

### Fks1-GFP localization is not altered during caspofungin treatment

To determine whether caspofungin treatment has a direct effect on the cellular localization of the β-1,3-glucan synthase, we examined the localization of the two components of the complex: Fks1 and Rho1, each as a fusion protein tagged with GFP. We treated *C. neoformans* strains expressing either endogenous Fks1-GFP or overexpressed GFP-Rho1 in a gradient of caspofungin concentrations and assessed localization over one hour of treatment (Figure S1). Both Fks1-GFP and GFP-Rho1 localized to endomembranes, including structures that resemble perinuclear endoplasmic reticulum staining (Pianalto *et al*. 2018). Similarly, both proteins localized to the outer edge of the cell. Fks1-GFP, in particular, appeared to localize to the plasma membrane, but in distinct patches rather than to the entire cell surface. These data support the hypothesis that this and other cell surface proteins are located to specific sites or areas at the cell surface (Simons and Toomre 2000; Malínská *et al*. 2003; Siafakas *et al*. 2006). We determined that there was no significant difference in Fks1-GFP or GFP-Rho1 localization in response to caspofungin treatment.

### Inhibition of efflux pumps increases potency of caspofungin

Recent studies suggest that caspofungin is not efficiently maintained at high intracellular concentrations in *C. neoformans* (Huang *et al*. 2016). We therefore hypothesized that this organism might be preventing intracellular accumulation of caspofungin through the activity of one or more efflux pumps. To address this hypothesis, we performed a drug interaction assay to determine whether caspofungin potency might be enhanced in the presence of clorgyline, a monoamine oxidase inhibitor that also acts as an inhibitor of ABC efflux pumps in other fungal species (Holmes *et al*. 2012). Indeed, we found that clorgyline and caspofungin had an FIC index ranging from 0.18 and 0.3125, indicating a strong synergistic interaction between caspofungin and clorgyline (Table 3). These data suggest a role for efflux pumps in the innate tolerance of *C. neoformans* to caspofungin treatment.

## DISCUSSION

Despite the medical importance of fungal infections, few novel antifungal drugs have been introduced in recent years. Echinocandins, one of the most recently approved classes of antifungal drugs, were especially promising given their limited toxicity profile and novel mechanism of action. However, these agents display limited efficacy against several fungal classes, including the thermally dimorphic fungi and *Cryptococcus* species (Abruzzo *et al*. 1997; Bartizal *et al*. 1997; Espinel-Ingroff 1998). Therefore, we strove to identify cellular processes that, when inhibited, might serve as potential targets for combinatorial therapy with echinocandins for treatment of cryptococcal infection.

Our screen, using a collection of *C. neoformans* mutant strains, revealed that cryptococcal tolerance to the echinocandin drug caspofungin is likely a multi-faceted phenomenon. We identified several pathways and processes that seem to be important for full tolerance of caspofungin, including some that have previously been associated with cell surface integrity, such as chitin biosynthesis and the phosphatase Ppg1, or general stress tolerance, such as calcineurin. Indeed, one of the most caspofungin-sensitive mutants that we identified was a *ppg1*Δ mutant. Interestingly, we did not see hypersensitivity among mutants in the cell wall integrity MAPK pathway, which suggests that non-canonical cell-wall integrity signaling could be playing a role in the response to caspofungin-induced cell wall stress (Gerik *et al*. 2005).

A recent study assaying *C. neoformans* deletion collections and random mutants for caspofungin sensitivity identified several other processes that were not identified in this study. Huang *et al*. determined that mutation of the Cdc50 lipid flippase β-subunit increased the sensitivity of this organism to caspofungin and fluconazole (Huang *et al*. 2016). By using a fluorescently labeled caspofungin molecule, they also determined that the *cdc50*Δ mutant strain displayed increased uptake of this drug compared to WT strains, suggesting that the altered membrane integrity and lipid content in the mutant strain might allow better penetration of the drug into the cell and, therefore, higher efficacy. This observation implies that overcoming the issue of limited entry into or accumulation of the drug in the cryptococcal cell might result in enhanced fungal cell killing. To study the importance of drug efflux, we assessed the effect of the efflux pump inhibitor clorgyline on caspofungin activity, and we found strong synergy between these two compounds. Together these data suggest that both poor entry of caspofungin into the cell and efflux of the drug from the cell could be preventing full caspofungin potency in *C. neoformans*.

In validation of our screening methods, we identified the calcineurin B subunit mutant as a hypersensitive strain. Calcineurin signaling is required for full virulence of *C. neoformans* and is involved in the response to elevated temperatures, as well as response to caspofungin and other cellular stresses (Odom *et al*. 1997; Cruz *et al*. 2000; Del Poeta *et al*. 2000; Fox *et al*. 2001). Similarly, calcineurin signaling has been shown to be required for paradoxical growth in the presence of high concentrations of caspofungin in *Candida* species and *Aspergillus fumigatus* (Walker *et al*. 2008; Fortwendel *et al*. 2010; Lee *et al*. 2011; Walker *et al*. 2012; Juvvadi *et al*. 2015). In *C. neoformans*, calcineurin is known to regulate one transcription factor, Crz1, and many calcineurin-dependent cellular processes are likely mediated by Crz1 (Lev *et al*. 2012). However, recent work demonstrated that calcineurin likely has multiple downstream effectors in addition to Crz1 (Chow *et al*. 2017). Our work suggests that the calcineurin-regulated response to caspofungin likely involves both the activity of the Crz1 transcription factor as well as Crz1-independent calcineurin targets. To begin to address which calcineurin-regulated target proteins might be working in conjunction with Crz1, we assessed the effect of mutants of several genes that are known to be dephosphorylated in a calcineurin-dependent manner (Park *et al*. 2016). However, none of these individual mutants revealed caspofungin hypersensitivity phenotypes. Future work could probe other calcineurin targets for roles in caspofungin tolerance in this organism.

We also probed multiple processes regulated by the Pfa4 palmitoyltransferase to explore whether the phenotype of the palmitoyltransferase mutant could be attributed to dysfunction of one or more of its downstream targets. Interestingly, we determined that the *C. neoformans* Ras1-mediated thermotolerance pathway is required for tolerance of caspofungin treatment in this organism. This signaling pathway responds to extracellular cues to control the activation of the Cdc42 GTPase, a protein that directs actin cytoskeleton polarization as well as directional growth and budding (Ballou *et al*. 2009). These changes in cellular structure appear to be mediated by septin proteins such as Cdc3 and Cdc12. In the absence of any of these pathway proteins, the cryptococcal cell is growth-impaired under many stressful conditions such as mammalian physiological temperatures (Nichols *et al*. 2007; Ballou *et al*. 2009, 2013a). This impairment is likely due to failed actin polarization resulting in defective budding and cell reproduction. Control of actin polarization is a documented response to cell wall stress in *S. cerevisiae*, required for the transient depolarization of Fks1 to deal with widespread cell wall damage (Delley and Hall 1999). Additionally, depolarization of the actin cytoskeleton has been associated with aberrant cell wall component deposition, likely due to the mislocalization of vesicles containing cell wall biosynthesis proteins such as chitin synthases and Fks1 (Gabriel and Kopecká 1995). Although mutants of two members of this pathway, *RAS1* and *CDC42*, were present in the collections tested in this study, their growth phenotypes are only apparent at elevated temperatures, and their caspofungin susceptibility would not have been identified at the lower temperatures tested.

Since Ras and Ras-related proteins are required for pathogenesis in many fungal systems, there have been concerted efforts to identify small molecules that inhibit Ras protein function. One of the most promising directions includes targeting protein prenyltransferases, enzymes that add lipid moieties to the C-terminal regions of Ras-like GTPases, directing the localization of these proteins to cellular membranes, their sites of function (Vallim *et al*. 2004; Nichols *et al*. 2009; Esher *et al*. 2016). Prenylation inhibitors are predicted to alter the localization and function of many cellular proteins in addition to Ras, potentially having broad antiproliferative effects in divergent cell types. Structural studies have identified fungal-specific features of farnesyltransferase enzymes, suggesting that antifungal specificity could be engineered into farnesyltransferase inhibitors (FTIs) (Hast *et al*. 2011; Mabanglo *et al*. 2014). Antifungal susceptibility testing demonstrated *in vitro* synergy between the activities of caspofungin and FTIs, the basis of which was identified in our screen. Importantly, the FTIs tested have limited efficacy themselves. However, as newer and more potent antifungal FTIs are identified, these agents might be used as adjunctive therapies to enhance the effect of caspofungin. It remains to be determined whether this synergy is due to FTI-mediated enhanced intracellular stability of echinocandins or synergistic activity in altered morphogenesis.

Interestingly, one of the most sensitive mutants identified in this study contained a deletion of the *C. neoformans MSH1* ortholog. However, this mutation was not directly responsible for the change in caspofungin sensitivity of the mutant strain. This gene has been previously described to play a role as a MutS homolog mismatch repair protein in the mitochondria in *S. cerevisiae* (Reenan and Kolodner 1992). We therefore hypothesized that this mutant strain might have accumulated additional mutations that were directly responsible for the caspofungin hypersensitivity phenotype. Indeed, mutations in mismatch repair proteins are associated with increased mutation rates, often associated with increased drug resistance (Healey *et al*. 2016; Billmyre *et al*. 2017). While this strain had few mutations in the nuclear genome, one of these was a nonsense mutation in a component of the COP9 signalosome which truncated the gene product. The COP9 signalosome has been implicated in fungal development, including involvement in the S-phase checkpoint in *S. pombe* as well as sexual development in *A. nidulans* (Mundt *et al*. 1999, 2002; Busch *et al*. 2003). This intracellular complex of proteins functions within the ubiquitin-proteasome pathway by controlling the activity of cullin-RING E3 ubiquitin ligases (Wei and Deng 2003). Our ability to complement the mutant strain with a WT copy of the *CSN4* gene, rather than the *MSH1* gene, strongly supports our hypothesis that COP9 signalosome complex dysfunction is the reason for this strain’s hypersusceptibility to caspofungin. The *CSN4STOP* strain displays a slight growth rate defect, which could suggest a potential cell cycle defect as seen in *S. pombe* mutants for COP9 signalosome components (Mundt *et al*. 1999).

We also determined that chitin biosynthesis is upregulated during caspofungin treatment, and the Chs3 chitin synthase is required for caspofungin tolerance. As caspofungin inhibits the biosynthesis of the cell wall component β-1,3-glucan, upregulation of other cell wall biosynthesis or cell wall-modifying enzymes may compensate for altered glucan content. Indeed, in *Aspergillus fumigatus*, cell wall chitin deposition, as well as the expression of chitin synthase genes, increases during caspofungin treatment (Fortwendel *et al*. 2009, 2010). Additionally, *Candida* clinical isolates with naturally increased levels of chitin, as well as strains grown in chitin-inducing conditions, demonstrate increased survival during caspofungin treatment (Walker *et al*. 2008, 2012; Lee *et al*. 2011). A similar phenomenon appears to be occurring in *C. neoformans:* during caspofungin treatment, there are increased levels of chitin both by CFW staining and by an *in vitro* biochemical quantification of chitin monomers. There is a similar increase in chitosan levels. As *Cryptococcus* species typically display higher levels of chitosan in the cell wall than other pathogenic fungal species (Banks *et al*. 2005), these inducible cell wall changes in chitin/chitosan content likely represent a conserved mechanism by which *C. neoformans* adapts to caspofungin treatment. Additionally, the increased baseline chitosan levels in the *C. neoformans* cell wall may also contribute to its innate tolerance to this drug.

In conclusion, we have used a genetic screen to identify *C. neoformans* strains with altered susceptibility to caspofungin to better understand the mechanism by which this species tolerates echinocandins. Several varied cellular processes were represented among the strains with enhanced caspofungin susceptibility. It is possible that some of these processes enhance the intracellular accumulation of the drug, helping to prevent the low concentrations of caspofungin in *C. neoformans* cells suggested in prior studies. As new fungal-specific inhibitors are developed for Ras protein localization and chito-oligomer synthesis, these agents may provide promising new directions for combination antifungal therapy.

## ACKNOWLEDGMENTS

This work was supported by NIH R01 grant AI074677 (J.A.A.) and P01 AI104533, as well as a National Defense Science and Engineering Grant to K.M.P. awarded by the Department of Defense, Office of Scientific Research, 32 CFR 168a. K.M.P. was also supported by a Duke University Bass Instructional Fellowship. The authors would like to thank the Madhani laboratory and NIH funding R01AI100272 for the deletion mutant collection in *C. neoformans*. Additionally, the authors would like to thank Joseph Heitman for support of and insight into this project.

